# Bioinformatic analyses of plasmid resistome changes in pOXA-48

**DOI:** 10.1101/2022.03.02.482687

**Authors:** Stephen Fordham, Anna Mantzouratou, Elizabeth Anne Sheridan

**Affiliations:** Bournemouth University, Department of Life & Environmental Sciences,, Talbot Campus Fern Barrow, Poole BH12 5BB; University Hospitals Dorset NHS Foundation Trust, Department of Medical Microbiology,, Poole Hospital, Longfleet road, BH15 2JB

**Keywords:** Klebsiella, plasmids, antibiotic resistance

## Abstract

Infections caused by carbapenem resistant Enterobacteriales (CPE) represent a significant threat in clinical settings. *bla*_OXA-48_ is one of the most frequent carbapenemase genes among Enterobacteriales. The *bla*_OXA-48_ is typically encoded on the prototypical IncL conjugative pOXA-48 plasmid. The pOXA-48 plasmid encodes only the *bla*_OXA-48_ resistance gene. However, aminoglycoside and extended spectrum β-lactamase (ESBL) resistance genes have also been detected on the same pOXA-48 plasmid backbone. These pOXA-48 plasmids encoding additional antimicrobial resistance (AMR) genes have been associated with both poor patient outcome and increased minimal inhibitory concentrations (MICs) to antibiotics including broad-spectrum cephalosporins.

The *bla*_OXA-48_ gene was sourced from the pOXA-48 reference plasmid and set as a query using the BLASTn tool. Non-duplicate *bla*_OXA-48_ containing plasmids were downloaded, incompatibility typed and annotated for resistance genes using ResFinder 4.0. Bioinformatic analyses identified three distinct variants of the pOXA-48 plasmid encoding 4, 5, and 6 antimicrobial resistance genes. All plasmids encoded the ESBL *bla*_CTX-M-14b_, *bla*_OXA-48_ and either 2, 3 or 4 aminoglycoside resistance genes, in addition to conjugative transfer machinery. Plasmid variants 1 and 3 encoded aminoglycoside genes bracketed between IS26 and ISEc63 insertion elements, forming a potential transposon. The potential transposon structure had resemblance to the Tn5393 transposon (accession: M96392), including both *aph(3’’)-Ib, aph(6)-Id* genes, and a Tn3 resolvase. The IS element ISEcp1 lies upstream of *bla*_CTX-M-14b_. All three plasmid variants appear related. Notably, all pOXA-48 plasmid variants were identified in multiple countries. In particular, variant 1 including 6 AMR genes was detected in 7 unique countries.

Plasmids encoding additional AMR genes were associated with clinical/surveillance samples suggesting antibiotic pressure in clinical settings may promote changes in the resistome of pOXA-48. Acquisition of pOXA-48 resistant plasmids carrying additional AMR genes beyond *bla*_OXA-48_ can change the resistome of susceptible isolates in a single-step, rendering previously susceptible strains refractory to almost all available treatment options.

## 1. Introduction

Mobile genetic elements (MGEs) promote the dissemination of carbapenem resistance genes. MGEs including plasmids and transposons help spread resistance determinants contributing to the rise in resistance to β-lactams, escalating clinical infections and poor clinical outcomes. *K. pneumoniae* and *Escherichia coli* (*E.coli*) are the dominant carbapenemase producing Enterobacterales (CPE) harboring *bla*_OXA_. The carbapenemase *bla*_OXA-48_ was first characterized in a *K. pnuemoniae* isolate from Turkey in 2001 (Poirel *et al*., 2004). *bla*_OXA-48_, the dominant *bla*_OXA_ variant identified in hospitals settings is now distributed worldwide. Commonly encoded on a pOXA-48 plasmid backbone, *bla*_OXA-48_ is typically bracketed by two identical insertion sequences IS*1999*, an IS4 family element involved in both the mobilization and expression of beta-lactam resistance genes. Together, the structure including a LysR transcriptional regulator form a composite transposon, Tn*1999* embedded within a *tir* transfer inhibition gene (Poirel *et al*., 2011).

In hospitals, *bla*_OXA-48_ disseminates via high-risk hospital adapted clones carrying the prototypical pOXA-48 plasmid. pOXA-48 spreads between hospitalized patients, colonizing their gut microbiota. Once colonized, conjugation mediates horizontal gene transfer (HGT) of the pOXA-48 plasmid to resident members of the gut microbiota. Conjugation assays reveal the IncL pOXA-48 plasmid exhibits efficient intraspecies and intergenus transconjugation (Hamprecht *et al*., 2019; León-Sampedro *et al*., 2021). Frequent plasmid transfer provides a test bench for new bacterium-pOXA-48 combinations. Acquisition of pOXA-48 by some of these bacteria results in low or even beneficial fitness costs, enabling the plasmid to persist and even disseminate to new hosts (Valle *et al*., 2020). pOXA-48 can persist in the human gut throughout hospital stays and can be detected months or even years later in subsequent admissions in different strains to the original colonizing strain (Pantel *et al*., 2016; Hernández-García *et al*., 2019; León-Sampedro *et al*., 2021).

Nosocomial outbreaks of OXA-48 producing *K. pneumoniae* have been associated with fatal outcomes. Invasive infections, notably bacteremia, caused by *K. pneumoniae* isolates carrying *bla*_OXA-48_, are associated with high-mortality (Navarro-San Francisco *et al*., 2013; Madueño *et al*., 2018; Pantel *et al*., 2016; Taoufik *et al*., 2019; Rodríguez *et al*., 2021). Further, carbapenem therapy in patients infected with *bla*_OXA-48_ has been linked with a high risk of treatment failure (Balkan *et al*., 2014; Shaidullina *et al*., 2020). Moreover, OXA-48 producing *K. pneumoniae* outbreaks are long-lasting and expensive to manage. In the UK, the estimated cost for managing an OXA-48 *K. pneumoniae* outbreak was approximately £400,000 (Lim *et al*., 2020).

OXA-48 carrying plasmids can be resistant to carbapenems but remain susceptible to cephalosporins unless additional beta-lactamase or other resistance genes are present. First line antimicrobial disk testing combinations for routine clinical samples can easily fail to detect the presence of the enzyme and the carbapenem-resistant cephalosporin susceptible phenotype is not confidently attributed to carbapenemase production using automated systems such as MicroScan NBC39 and Vitek 2 in diagnostic laboratories (Woodford *et al*., 2010). OXA-48 producers can therefore disseminate widely undetected. Two further factors additionally complicate OXA-48 producer detection in clinical settings. First, OXA-48 produces can often have low minimum inhibitory concentrations (MICs), below clinical breakpoints to both imipenem and meropenem (Dimou et al., 2012, PHE, 2016). Second, due to the low ratio of infection to colonization, clinical specimens serve as an inaccurate and late identifier for the presence of OXA-48 producers in a clinical setting (Lim *et al*., 2020). These factors combined, make the detection of CPE harboring *bla*_OXA-48_ challenging. Pervasive OXA-48 producers in clinical settings may engender a scenario whereby the resistome of pOXA-48 plasmids have opportunities to change, potentially requiring more resistance genes.

The pOXA-48-like plasmid typically only encodes the *bla*_OXA-48_ gene. The presence of additional AMR genes on the pOXA-48 backbone has been reported. For example, 2 enterobacterial strains, *E. coli* ESS-1 and *Enterobacter cloacae* ESS-2 carried the ESBL *bla*_CTX-M-15_ gene on a pOXA-48 backbone. The strains encoding *bla*_CTX-M-15_ displayed resistance to broad-spectrum cephalosporins in a patient who had recently been hospitalized in Algeria (Potron *et al*., 2013). Worryingly, fatality was reported in patients infected with a *K. pneumoniae* strain carrying a pOXA-48-like plasmid pJEG011 encoding aminoglycoside resistance genes, the ESBL *bla*_CTX-M-14_, and *bla*_OXA-48_. Strain Kp001 harboring pJEG011 was resistant to aminoglycosides, and β-lactams, including meropenem (Espedido *et al*., 2013). Moreover, in Palestine, the Netherlands and China, pOXA-48 plasmids encoding both aminoglycoside resistance genes and *bla*_CTX-M-14_ have been reported (Chen *et al*., 2015; Hendrickx *et al*., 2020; Wang *et al*., 2021).

The acquisition of additional AMR genes on the single broad host conjugative plasmid, pOXA-48, has the potential to immediately change the resistome of recipient strains in a single-step process, rendering previously susceptible strains refractory to almost all available antibiotics. *In-silico* antibiotic resistance gene analyses was undertaken to determine resistome patterns in pOXA-48 like plasmids sourced from NCBI using bioinformatics analyses. Specifically, we aimed to detect common AMR genes on the pOXA-48 backbone, determine their genomic architecture, relatedness, presence of conjugation genes and international distribution.

## 2. Method

### 2.1. Plasmid Sample Acquisition

Plasmid contigs encoding the *bla*_OXA-48_ allele were sourced from the National Center for Biotechnology Information (NCBI). The nucleotide reference sequence of *bla*_OXA-48_ was derived from reference pOXA-48 plasmid JN626286.1 (Poirel *et al*., 2011). To retrieve pOXA-48 like plasmids encoding *bla*_OXA-48_, the BLASTn program from NCBI was used. To retrieve all plasmid sequences encoding *bla*_OXA-48_, the max target sequences parameter on NCBI BLASTn was adjusted from the default of 100 to 1000. The resultant hit description table from NCBI was downloaded. Non-duplicate plasmid hits were sourced as of 16/12/21. Plasmid samples were sourced from the description table by filtering for ‘plasmid’ using the Python library Pandas. Additionally, strict parameters were applied to retrieve pOXA-48-like plasmids. For all plasmid sequences, a 100% query coverage and nucleotide identity for the gene encoding *bla*_OXA-48_ were required. The strict nucleotide identity criterion was chosen to acquire plasmids encoding *bla*_OXA-48_. BLASTn hits with nucleotide identity of < 99.9% return different *bla*_OXA_ alleles. Finally, to retrieve pOXA-48 like plasmids, a size criterion between 60-75 kb for the plasmids was applied. This criterion was chosen to remove chromosomal hits, short contigs encoding *bla*_OXA-48_, and to capture the variation in size of pOXA-48 carrying plasmids reported from 60 kb (Poirel *et al*., 2011; Findlay *et al*., 2017) up to 75 kb (Hamprecht *et al*., 2019). Together, the filtering criteria returned 196 plasmids including reference pOXA-48 plasmid JN626286.1, with a size range of 60-75 kb, all encoding the 798 bp *bla*_OXA-48_ gene. Plasmid acquisition is summarized in workflow Figure 1A.

**Figure 1:**
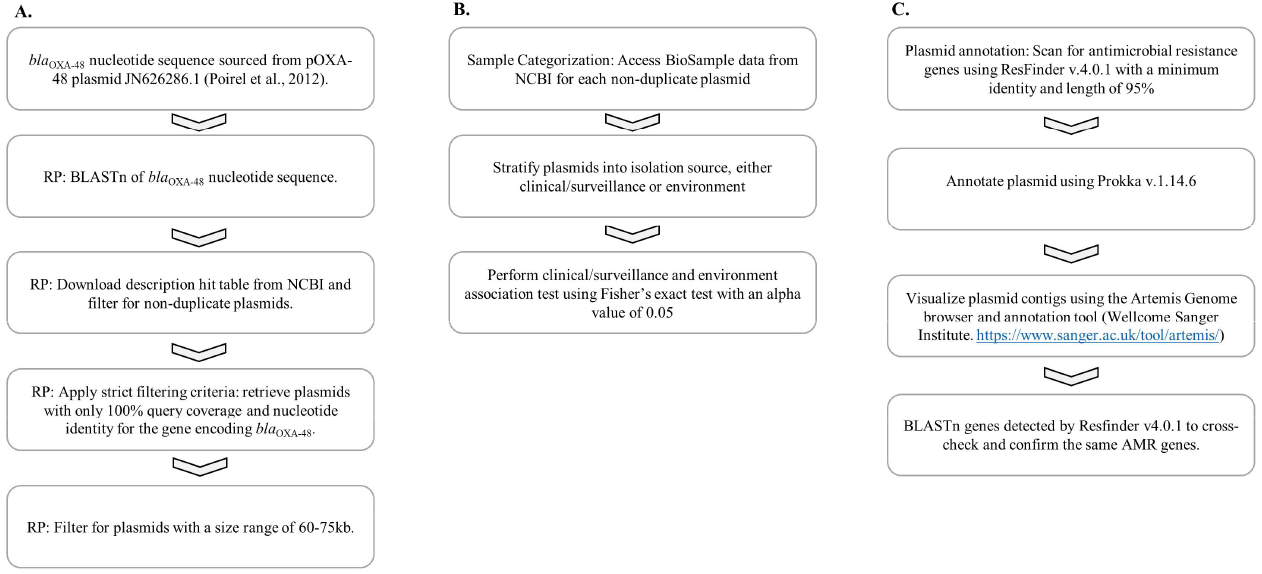
Method Workflow. A Retrieve plasmids (RP). B. Categorize plasmids. C. Annotate and visualize plasmids.

The pOXA-48-like plasmids were stratified into one of two categories; clinical/surveillance or environment based on the isolation source listed on the Biosample data for each plasmid sample, see Supplementary Table S1.

### 2.2. Plasmid Annotation

Plasmid samples were annotated to identify resistance genes and determine the genomic context of those resistance genes. FASTA files were annotated using PROKKA version 1.14.6 using default parameters (Seemann *et al*., 2014). FASTA contigs were incompatibility (Inc) typed using PlasmidFinder using a minimum identity and length of 98% (Carattoli and Hasman, 2019). Each plasmid contig was also scanned for resistance genes using ResFinder v4.0 with a minimum identity and length of 95% (Zankari *et al*., 2012).

To help confirm AMR gene annotations, a small validation using a subset of plasmids encoding either the ESBL *bla*_CTX-M-14_, aminoglycoside resistance genes, and the *bla*_OXA-48_ carbapenemase gene was performed. Detected AMR genes confirmed by PCR/sequencing coupled with MIC testing from reported studies was compared to predictions made using ResFinder v.4.0 (Table A1). Furthermore, AMR genes were also annotated using Prokka v.1.14.6, visualised using Artemis, and Blasted using NCBI to confirm their presence. The genomic architecture of the pOXA-48 variants encoding AMR genes were visualized using Easyfig version 2.2.2 (Sullivan *et al*., 2011). The plasmid acquisition process is shown in Figure 1C.

## 3. Results

### 3.1. Plasmid Source and Size

*K. pneumoniae* was the most abundant species harboring pOXA-48-like plasmids. *K. pneumoniae* carried 71.94% (*n*=141/196) of pOXA-48-like plasmids, 3.81-fold higher than the next most abundant pOXA-48-like carrying species, *E.coli* (Figure 2a). Combined, both *K. pneumoniae* (*n*=141/196) and *E.coli* (*n*=37/196) made up 90.8% of the species carrying pOXA-48-like plasmids. The pOXA-48 plasmid was detected in 11 additional species (Figure 2a). These include *Enterobacter cloacae* (*n*=4), *Citrobacter freundii* (*n*=4), *Enterobacter hormaechei (*n*=2), Klebsiella aerogenes* (*n*=1), *Raoultella ornithinolytica* (*n*=1), *Pluralibacter gergoviae* (*n*=1), *Citrobacter werkmanii* (*n*=1), *Enterobacter roggenkampii* (*n*=1), *Raoultella planticola* (*n*=1), *Serratia marcescens* (*n*=1) and *Proteus mirabilis* (*n*=1). Across the 196 samples, 79.6% (*n*=156/196) of plasmid samples were derived from either clinical or surveillance samples, while 12.78% (*n*=25/196) of samples were sourced from the environment (Figure 2b). All plasmids were typed as IncL/M plasmids.

**Figure 2.**
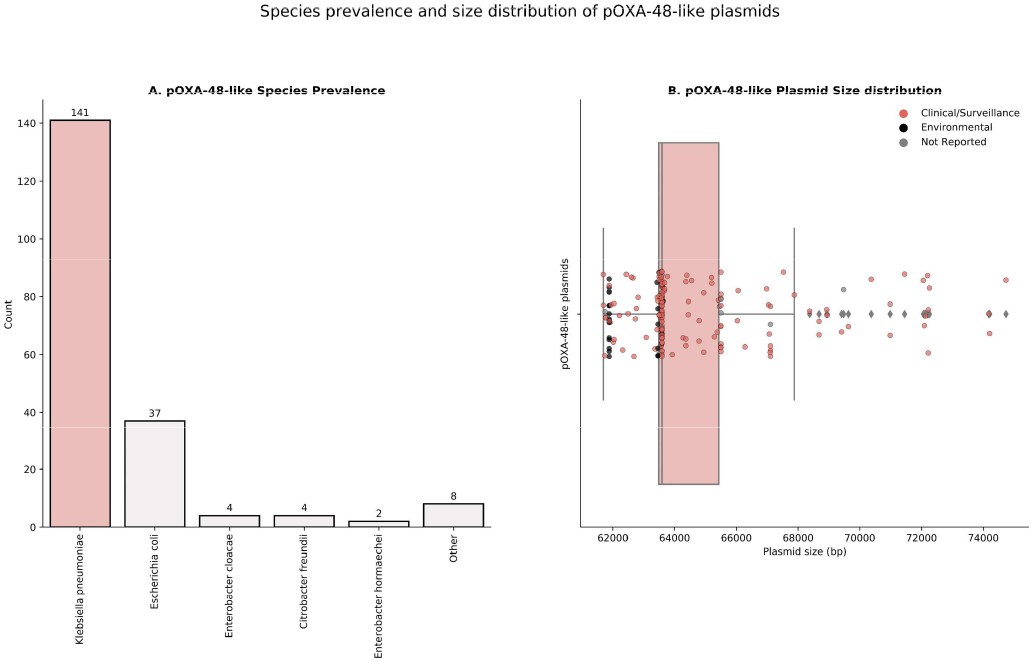
Species prevalence, source, and size distribution of all pOXA-48 plasmid samples. A. pOXA-48-like plasmids were identified in 13 unique species. *K. pneumoniae* (*n*=141/196) and *E.coli* (*n*=37/196) made up 90.8% of the species carrying pOXA-48-like plasmids. B. Plasmid samples were most commonly identified from clinical or surveillance sources, 79.6% (*n*=156/196) versus environmental sources 12.78% (*n*=25/196). The 34 pOXA-48 plasmids carrying multiple AMR genes have a larger plasmid size, and a more divergent nucleotide identity and coverage, relative to the reference plasmid, pOXA-48 (accession JN626286.1).

The median plasmid size was 63589 bp, mean 64592 bp (standard deviation 2728 bp). The majority of pOXA-48-like plasmids only encoded *bla*_OXA-48_ (*n*=162/196). Despite this, 34 pOXA-48 encoding plasmids carried more than one antimicrobial resistant (AMR) gene. Plasmids encoding multiple AMR genes had a larger mean (69433 bp versus 63576 bp) and median size (68937 bp versus 63539 bp) relative to the reference pOXA-48 plasmid JN626286.1 (Poirel *et al*., 2011).

The pOXA-48-like plasmid was identified in 25 environmental samples (Supplementary Table S1). These samples have a close nucleotide coverage (mean=98.48%, median=100%, standard deviation=2.0%) and average nucleotide identity (mean=99.76%, median=99.79%, standard deviation=0.105%) against the reference pOXA-48 plasmid JN626286.1. Concerningly, 4 plasmid samples were identified from human food consumption sources, 2 from retail meats from China, 1 from cow milk in France and 1 from vegetables.

### 3.2. Antimicrobial resistance genes on pOXA-48 plasmids

A total of 9 AMR genes were detected from the 34 pOXA-48-like AMR plasmid samples. In addition to *bla*_OXA-48_ present in all plasmid samples, a further 8 AMR genes were detected. These include: *aph(3”)-Ib, aph(3’)-Vib, aph(6)-Id, bla*_CTX-M-14_, *bla*_CTX-M-14b_, *bla*_TEM-1B_, *qnrB*, and *qnrS1* encoding a resistance phenotype to the following antibiotics: kanamycin, ampicillin, ceftriaxone, meropenem, and ciprofloxacin. The occurrence of *bla*_OXA-48_ plus additional AMR gene(s) is associated with samples sourced from clinical/surveillance settings (Fisher’s exact test, *p* value = 0.009008).

Across the 34 plasmid samples encoding AMR genes, 3 common AMR variants were detected (Figure 3). The first variant, named pOXA-48 variant-1 included 9 samples which encoded the same 6 AMR genes. These included: *aph(3”)-Ib* (2), *aph(6)-Id, aph(3’)-Vib*, *bla*_CTX-M-14b_, and *bla*_OXA-48_. Interestingly, pOXA-48 variant-1 was detected from 7 independent countries, including Germany (*n*=2), China (*n*=2), UK (*n*=1), USA (*n*=1), Palestine (*n*=1), Australia (*n*=1), and the Netherlands (*n*=1). All pOXA-48 variant-1 samples were derived from clinical/surveillance sources (Supplementary Table S1).

**Figure 3.**
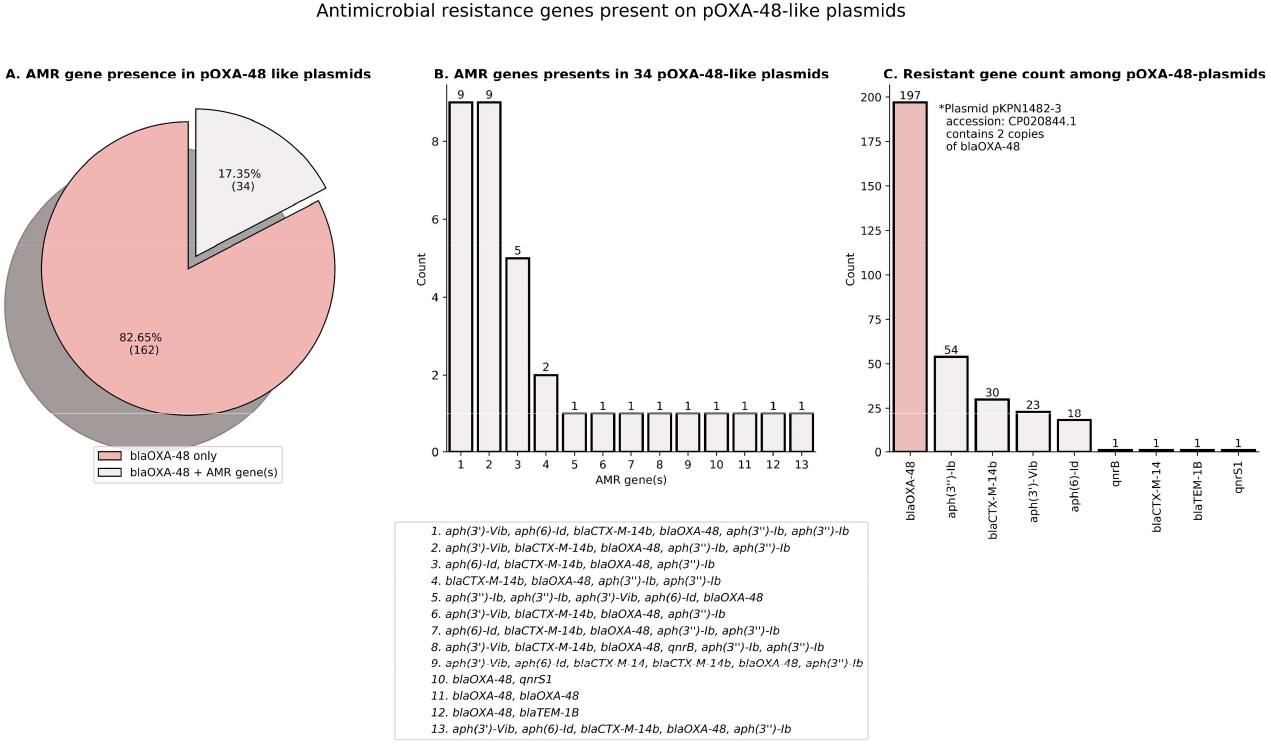
Antimicrobial resistance gene (AMR) distribution among pOXA-48 plasmids. A. From the sample pool of 196 pOXA-48 plasmids, 34 harbor AMR gene(s) in addition to *bla*_OXA-48_. B. The 3 most common variants encoded 9, 9, and 5 AMR genes, respectively. These included *aph(3”)-Ib* (2), *aph(6)-Id, aph(3’)-Vib*, *bla*_CTX-M-14b_, and *bla*_OXA-48_ (Key 1 in legend), *aph(3’’)-Ib* (2), *aph(3’)-Vib*, *bla*_CTX-M-14b_ and *bla*_OXA-48_ (Key 2 in legend), and *aph(6)-Id*, *bla*_CTX-M-14b_, *bla*_OXA-48_, *aph(3”)-Ib* (Key 3 in legend). C. A total of 9 AMR genes were detected in addition to the *bla*_OXA-48_ gene present in the Tn1999 transposon of pOXA-48-like plasmids. These included: *aph(3”)-Ib* (*n*=54), *bla*_CTX-M-14b_ (*n*=30), *aph(3’)-VIb* (*n*=23), *aph(6)-Id*(*n*=18), *bla*_TEM-1B_ (*n*=1), *qnrS1* (*n*=1), *bla*_CTX-M-14_ (*n*=1), and *qnrB* (*n*=1). 2 copies of the beta lactamase gene, *bla*_OXA-48_ were present in plasmid pKPN1482-3 (accession: CP020844.1). The nucleotide sequence of *bla*_CTX-M-14b_ differs from *bla*_CTX-M-14_ by one nucleotide substitution in the stop codon, TAA to TGA; both genes encode the same protein sequence.

The second variant, labelled pOXA-48 variant-2 included 9 samples encoding the same 5 AMR genes including *aph(3”)-Ib* (2), *aph(3’)-Vib*, *bla*_CTX-M-14b_ and *bla*_OXA-48_. pOXA variant-2 was sourced from 4 countries including Netherlands (*n*=4), Germany (*n*=2), Switzerland (*n*=2), and China (*n*=1). All samples, with the exception of one from Switzerland (plasmid accession CP083013.1) were derived from clinical/surveillance sources (Supplementary Table S1).

The third variant encoded the same 4 AMR genes: *aph(3”)-Ib, aph(6)-Id*, *bla*_CTX-M-14b_, and *bla*_OXA-48_. pOXA-48 variant 3 was sourced from just 2 countries including Saudi Arabia (*n*=1), and the Netherlands (*n*=4). All pOXA-48 variant 3 samples were isolated from clinical/surveillance samples (Supplementary Table S1).

### 3.3. pOXA-48 Variant 1

pOXA-48 variant 1 includes 2 insertions relative to the reference plasmid, pOXA-48 (accession JN626286.1). The first insertion is a 3,095 bp fragment encoding the IS element ISEcp1 encoding a transposase, a gene encoding a hypothetical protein of 40 amino acids and the beta lactamase gene, *bla*_CTX-M-14b_. The ISEcp1-*bla*_CTX-M-14b_ fragment is inserted downstream of the z1226 gene in the intergenic region. The nested position of the insert for representative pOXA-48 plasmid variant 1, *K. pneumoniae* plasmid pIncL_M_DHQ1400954 (accession: CP016927.1) is shown in Figure 4, between 2 genes shaded dark gray: the z1266 gene and a gene encoding a hypothetical protein.

**Figure 4.**
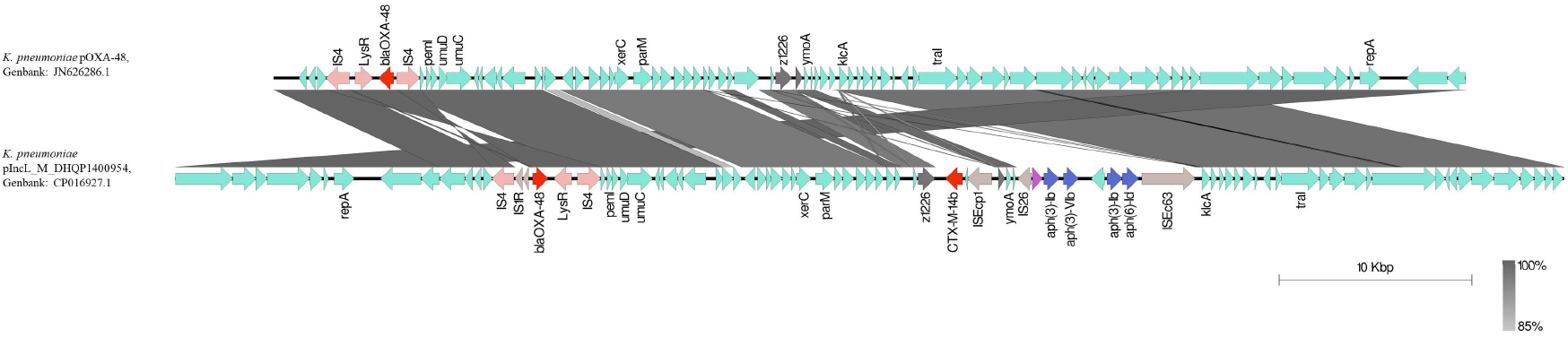
*K. pneumoniae* plasmid pIncL_M_DHQ1400954 (accession: CP016927.1), a representative member of pOXA-48 variant 1 carries two inserts relative to the reference plasmid pOXA-48 (accession JN626286.1). The first insertion is a 3,095 bp fragment encoding the ISEcp1 transposase, a gene encoding a hypothetical protein of 40 amino acids and the beta lactamase gene, *bla*_CTX-M-14b_. The ISEcp1-*bla*_CTX-M-14b_ fragment is inserted 37 nucleotides downstream of the z1226 gene in the intergenic region. The IS26-ISEc63 AMR fragment forms another independent insertion, positioned downstream of a hypothetical gene, after *ymoA*, at position 73 of the intergenic region, and upstream of *klcA* at position −201 in the intergenic region.

The second insert encodes 4 AMR genes, a ISPa14 transposase and a resolvase bracketed between two transposases: the IS26 transposase upstream, running in the opposite direction with respect to the AMR genes, and an ISEc63 transposase downstream. This insert fragment bears resemblance to the transposon Tn5393 (accession: M96392), where it includes the same *aph(6)-Id* gene, encoding streptomycin phosphotransferase. Upstream of the *aph(6)-Id* gene, 2 further AMR genes are found; *aph(3’’)-Ib* and *aph(3’)-Vib*, followed by the same shared *aph(3’’)-Ib* gene and Tn3 resolvase genes found in the Tn5393 transposon. Combined, the 4 AMR genes, alongside the resolvase, bracketed by 2 transposases may form a potential composite transposon, which include modules belonging to the Tn5393 transposon. The potential transposon forms a 9,292 bp fragment. The fragment includes the inverted repeat left (IRL) of ISEc63 and the inverted repeat right (IRR) of the IS26. The IS26-ISEc63 AMR bearing fragment positioning for representative pOXA-48 plasmid variant 1, *K. pneumoniae* plasmid pIncL_M_DHQ1400954 is shown in Figure 4.

9 samples harbored the 6 AMR genes common to pOXA-48 variant 1. 4 of these plasmid samples contained the complete structure of the potential composite transposon bracketed by IS26 and ISEc63 transposases. These included plasmids pIncL_M_DHQ1400954, pOXA-48-Pm, pJEG011, and pOXA-48-IR1251. Another 4 samples contained structures similar to the potential transposon. A comparison between 8 of these samples is shown in Figure 5.

**Figure 5.**
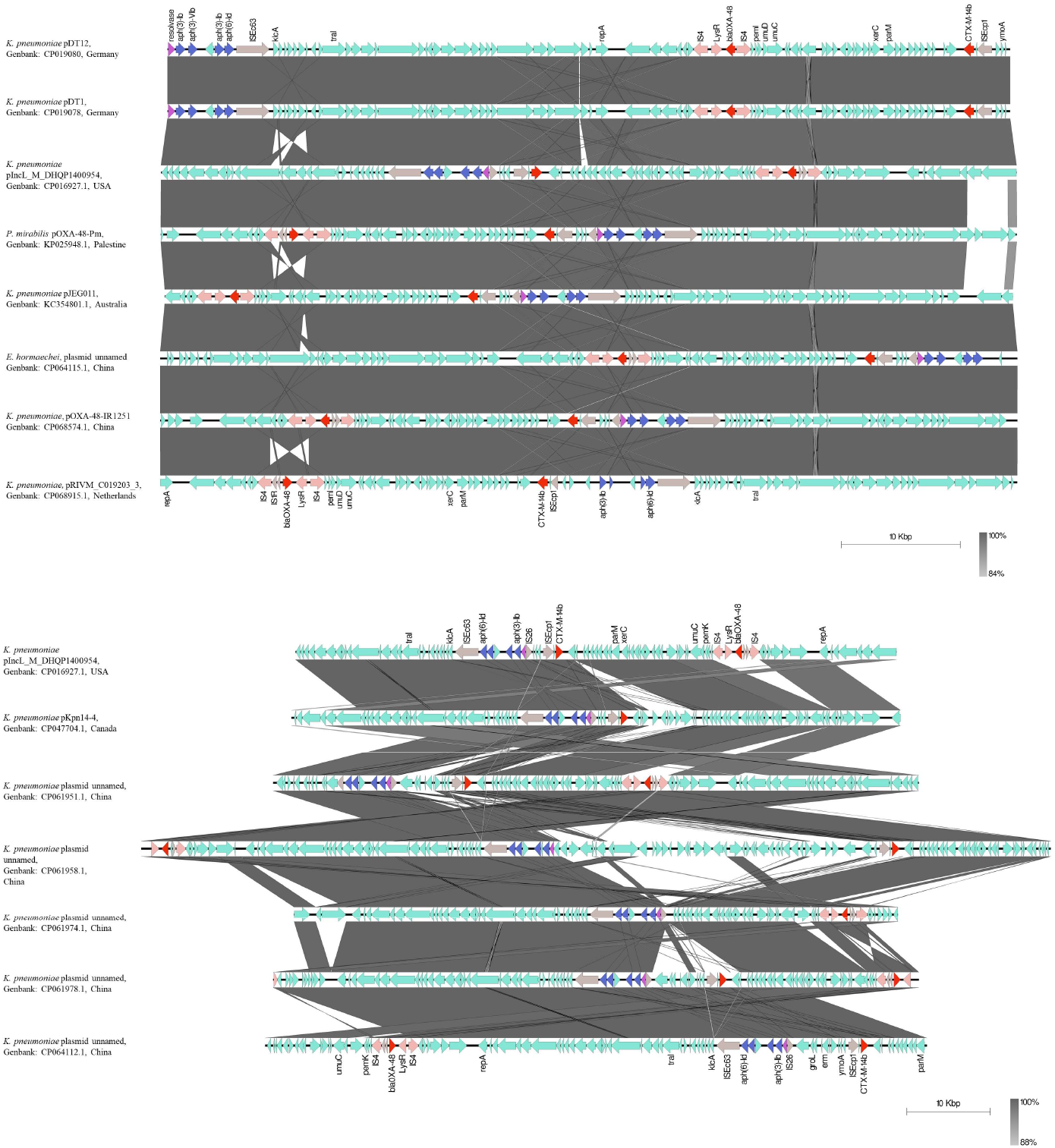
Genomic architecture of the potential composite transposon bracketed by insertion sequences IS26 and ISEc63. Top. The structure of the potential transposon carrying 4 AMR genes and a resolvase in shown in the pOXA-48-like plasmids. Bottom. The 9,292 bp potential transposon fragment was used as a search query for the NCBI program BLASTn. 6 plasmid hits are shown relative to representative plasmid pIncL_M_DHQP1400954 encoding the transposon structure.

In order to decipher the prevalence of the potential transposon, the BLASTn program from NCBI was used with the 9,292 bp fragment as the search query. In total, 15 plasmid hits were returned with coverage 100% (*n*=12), 99% (*n*=3), and percentage identity 100% (*n*=11), and >99.5% (*n*=4). The potential transposon was detected in 7 unique countries, including China (*n*=8), Netherlands (*n*=2), Palestine (*n*=1), the United Kingdom (*n*=1), Australia (*n*=1), USA (*n*=1), and Canada (*n*=1). 7 plasmid samples from the original 9 containing the 6 AMR genes common to pOXA-48 variant 1 were included in these 15 BLASTn hits.

Among the 9 samples carrying the 6 AMR genes common to pOXA-48 variant 1, the plasmids appeared related (Table 1). The additional plasmid hits, samples 10-17, which carried the 4 AMR genes in the IS26-ISEc63 fragment had high identity, ≥ 99.29% (*n*=8) and coverage ≥92% (*n*=6) relative to pOXA48-IR1251 was observed (Table 1). Collectively, these results confirm the similarity of the plasmids.

**Table 1.** pOXA-48 variant 1 relatedness.

| Sample | Plasmid | ST | Country | *Plasmid Size/Species | Accession | IS26-ISEc63 AMR fragment | bla<sub>CTX-M-14b</sub> | SNPs from plasmid pOXA48-IR1251) | Additional Notes | Identity to pOXA48-IR1251 | Coverage against pOXA48-IR1251 | Collection date |
| --- | --- | --- | --- | --- | --- | --- | --- | --- | --- | --- | --- | --- |
| 1 | pKpvST383L_2 | 383 | UK | 72057 bp Kp | CP034202.2 | Y | Y | 15 |  | 99.96% | 100% | 2018-04-12 |
| 2 | pDT1 | 383 | Germany | 70987 bp Kp | CP019078.1 | N | Y | 1 | No IS26 transposase | 100% | 99% | 2015-10-22 |
| 3 | pDT12 | 383 | Germany | 70987 bp Kp | CP019080.1 | N | Y | 5 | No IS26 transposase | 99.99% | 99% | 2016-03-29 |
| 4 | pIncL_M_DHQP1400954 | Not listed | USA | 72093 bp Kp | CP016927.1 | Y | Y | 284 | CP068574.1 has 1 6491 insertion | 99.58% | 100% | 2014-03-08 |
| 5 | pOXA48-Pm | Not listed | Palestine | 72127 bp Pm | KP025948.1 | Y | Y | 466 | CP068574.1 has 3 insertions totalling 9982 bp | 99.58% | 95% |  |
| 6 | pJEG011 | 101 | Australia | 71446 bp Kp | KC354801.1 | Y | Y | 2 | CP068574.1 has 1 777 bp insertion | 99.99% | 100% | 2013 |
| 7 | IR5283 | 418 | China | 72217 bp Eh | CP064115.1 | N | Y | 1 |  | 99.99% | 100% | 2015-08-21 |
| 8 | pOXA48-IR1251 | 11 | China | 72218 bp Kp | CP068574.1 | Y | Y | N/A | N/A | N/A | N/A | 2015-04-23 |
| 9 | pRIVM_C019203_3 | 147 | Netherlands | 72194 bp Kp | CP068915.1 | N | Y | 6 |  | 99.92% | 100% | 2019 |
| 10 | IR5654 | 11 | China | 67524 bp Kp | CP061956.1 | Y | N | 2 | CP068574.1 has 5 insertions totalling 4700 bp | 99.99% | 100% | 2015-11-01 |
| 11 | IR5392 | 34 | China | 79397 bp Ko | CP064112.1 | Y | Y | 5 | CP064112.1 has 2 insertions totalling 3584 bp | 100% | 92% | 2015-08-20 |
| 12 | IR5086 | 147 | China | 77479 bp Kp | CP061978.1 | Y | Y | 1 | CP061978.1 have 1 insertion totalling 5260 bp | 100% | 94% | 2014-02-11 |
| 13 | IR5726 | 11 | China | 109135 bp Kp | CP061958.1 | Y | Y | 1 | CP064112.1 has 8 insertions totalling 37,475 bp | 100% | 68% | 2017-08-09 |
| 14 | IR5755 | 1128 | China | 72450 bp Kp | CP061974.1 | Y | N | 1 | CP068574.1 has 5 insertions totalling 5339 bp, CP061974.1 has 2 insertions totalling 5578 bp | 100% | 92% | 2018-04-13 |
| 15 | IR5065 | 383 | China | 77477 bp Kp | CP061951.1 | Y | Y | 1 | CP061951.1 has 1 insertion totalling 5260 bp | 100% | 94% | 2013-04-29 |
| 16 | **pKpn14-4 | 376 | Canada | 73052 bp Kp | CP047704.1 | Y | Y | 676 | CP068574.1 has 4 15591 bp insertion, CP047704.1 has 5 insertions totalling 10049 bp | 99.29% | 87% | 2014-09-29 |
| 17 | pRIVM_C016345_2 | 383 | Netherlands | 75854 bp Kp | CP068304.1 | Y | Y | 3 | CP068304.1 has 1 insertion totalling 3669 bp | 99.9% | 95% | 2018 |

\*Kp: *Klebsiella pneumoniae*, Pm: *Proteus mirabilis*, Eh: *Enterobacter hormaechei*, Ko: *Klebsiella oxytoca*. Grey section denotes plasmids containing all 6 AMR genes common to pOXA-48 variant 1, white section includes samples with the IS26-ISEc63 AMR fragment including samples with/without CTX/OXA-48. \*\*no bla<sub>OXA-48</sub> present.

In China, the plasmids appeared related. Plasmid pOXA48-IR1251 was between 1-5 SNPs between the other 7 plasmids. All plasmids from China were sourced from K. *pneumoniae* isolates with the exception of one plasmid identified from *Enterobacter hormaechei* strain IR5283. The plasmid from the E. *hormaechei* was 1 SNP different from the plasmid pOXA48-IR1251 suggestive of recent horizontal cross-species transfer. Two plasmids isolated from *K. pneumoniae* strains IR5726, and IR5755 did include 8 insertions totaling 37,475 bp, and 2 insertions totaling 5,578 bp, however these were only 1 SNP different from plasmid pOXA48-IR1251.

Similarly, the plasmids from the Australia (pJEG011), the Netherlands (pRIVM_C016345_2 and pRIVM_C019203_3) and the UK (pKpvST383L_2) were 2, 3,6 and 15 SNPs different from plasmid pOXA48-IR1251 from China. Together, these results indicate the spread of a plasmid carrying multiple AMR genes.

In contrast, the plasmid from USA (pIncL_M_DHQP1400954), Palestine (pOXA-48-pm), and Canada (pKpn14-4) were 284, 466, and 676 SNPs different from the plasmid from China, pOXA48-IR1251. Furthermore, the plasmids from the USA (pIncL_M_DHQP1400954) and Canada (pKpn14-4) were divergent; 397 SNPs, and 5 insertions, totaling 10,049 bp. The plasmids from Palestine (pOXA-48-pm) the USA (pIncL_M_DHQP1400954)/Canada (pKpn14-4) were also divergent; 187 SNPs with a single insertion of 3527 bp, and 198 SNPs with 4 insertions totaling 6,588 bp, respectively. Taken together, the IS26-ISEc63 potential transposon may disseminate AMR genes by horizontal gene transfer (HGT) and independent import into a pOXA-48 IncL plasmid backbone.

#### 3.3.1 *Tra* Transfer Operon for pOXA-48 variant 1

The presence of the Tra transfer operon, HIJK*pri*LMNOPQRUWXY across the 9 samples bearing the 6 AMR genes common to pOXA-48 variant 1 was investigated. The nucleotide sequence of the *tra* transfer operon, responsible for plasmid conjugation was obtained from reference pOXA-48 plasmid JN626286.1 (Poirel *et al*., 2011). The *tra* operon encodes a sequence of 19,640 bp including the start of the operon, *traH* through to the end of the operon encoding *traY*. All samples had a nucleotide coverage of 100% and a nucleotide identity ≥ 99.83% with the exception of two plasmid samples. Plasmid sample, Pm-Oxa48 (accession: KP025948.1), sourced from a multi-drug resistant *Proteus mirabilis* strain lacked both *traX* and *traY* at the end of the operon, with only 85% coverage and average nucleotide identity of 97.60%, relative to the complete *tra* operon, HIJK*pri*LMNOPQRUWXY. Despite this, a transconjugation assay into recipient *E.coli* J53 cells revealed Pm-Oxa48 is a conjugative plasmid capable of transferring AMR genes co-residing on the same plasmid vector (Chen *et al*., 2015).

### 3.4. pOXA-48 Variant 2

pOXA-48 variant 2 encodes 5 AMR genes; *bla*_CTX-M-14b_, *bla*_OXA-48_, *aph(3”)-Ib* (2) and *aph(3’)-Vib*. Two copies of the *aph(3”-Ib)* gene, and *aph(3’)-Vib* alongside a recombinase are bracketed between 2 IS26 insertion transposases, forming a fragment of 6,660bp. pOXA-48 variant 2 is characterized by a series of insertion sequences and recombinase inserts located in close proximity. The IS26 AMR bracketed fragment, alongside insertion sequences is shown for representative pOXA-48 variant plasmid, p2-BD-13-OXA48 relative to the reference plasmid, pOXA-48 in Figure 6.

**Figure 6.**
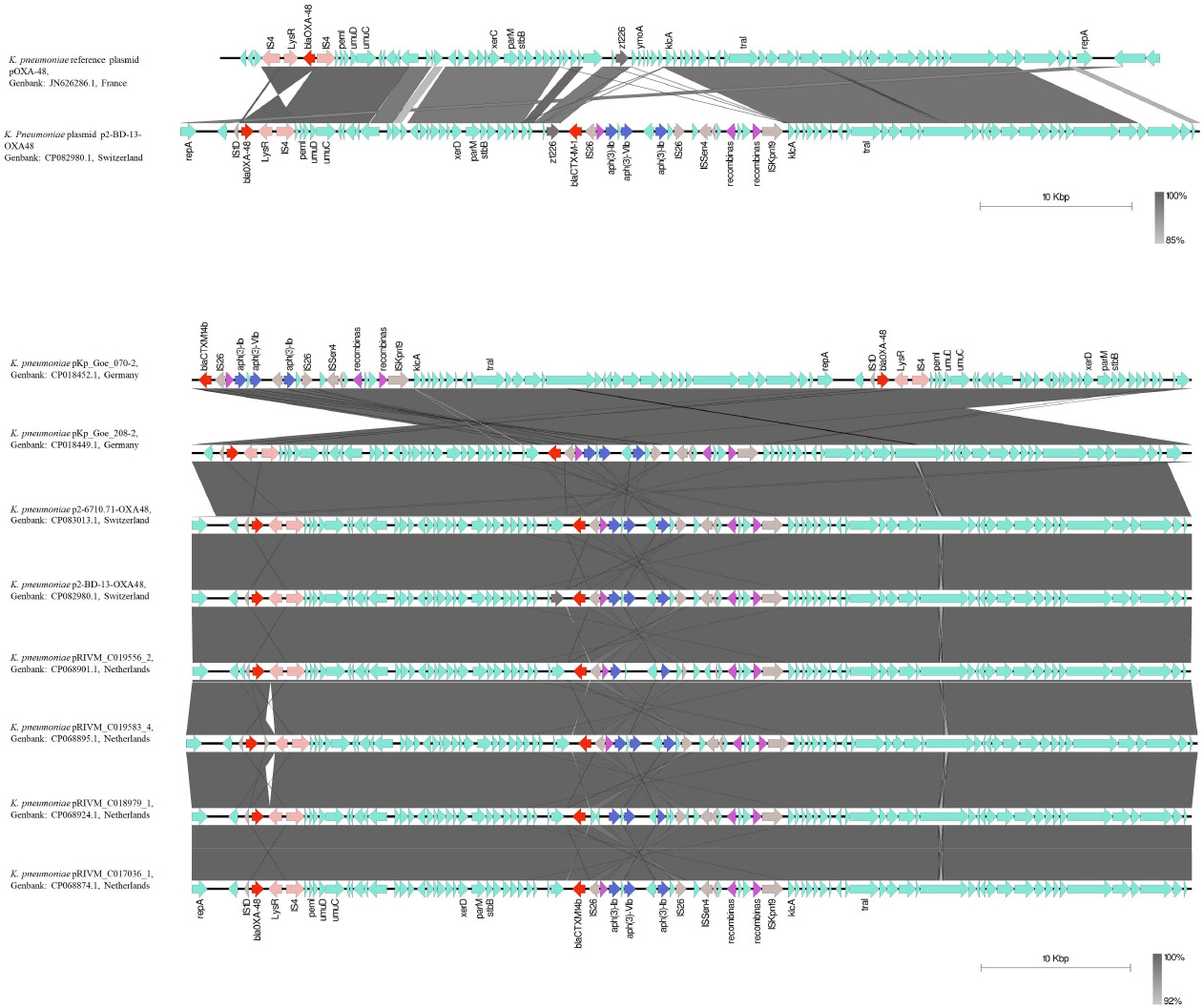
Genomic architecture of pOXA-48 variant 2. 3 AMR genes in genetic linkage with the same direction of transcription are bracketed between 2 IS26 insertion transposases, forming a fragment of 6,660bp. The fragment includes 2 copies of *aph(3’’-Ib)* (*strA*) gene, and *aph(3’)-VIb* alongside a resolvase/recombinase.

The plasmids encoding the 5 AMR genes were closely related. All 9 plasmids, with the exception of, p2-0113481141-OXA48 (accession: CP083077.1) had 100% coverage and ≥99.75% identity with plasmid pRIVM_C018979_1 (accession: CP068874.1) from the Netherlands. In fact, one plasmid from Switzerland and 2 from Germany had 100% coverage and 99.99% identity. 4 of 6 samples from the Netherlands were identified from the high-risk *K. pneumoniae* ST101.

Despite the international distribution, the plasmids are similar, whereby the 2 plasmids from Germany, pKp_Goe_070-2 (accession: CP018452.1), and pKp_Goe_208-2 (accession: CP018449.1) are only 8 and 9 SNPs different from plasmid pRIVM_C018979_1 (accession: CP068874.1) detected from a *K. pneumoniae* isolate from the Netherlands, and only 1 SNP different from one another. All isolates from the Netherlands are between 11-43 SNPs from pRIVM_C018979_1. Furthermore, plasmid p2-BD-13-OX48 detected from Switzerland (accession: CP082980.1) is 8 SNPs different from pRIVM_C018979_1. The second plasmid sample from Switzerland, p2-0113481141-OXA48 (accession: CP083077.1) is 128, and 125 SNPs different from pRIVM_C018979_1 and the p2-BD-13-OX48, respectively, suggesting plasmid p2-0113481141-OXA48 may be more distantly related to the other 9 samples.

#### 3.4.1 *Tra* Transfer Operon for pOXA-48 variant 2

Supporting the close similarity between the plasmids, all samples had 85% coverage and 97.60% identity against the reference *tra* operon, HIJK*pri*LMNOPQRUWXY from pOXA-48 plasmid JN626286.1 (Poirel *et al*., 2011). The plasmids samples with the AMR genes between the IS26 insertion sequences, all lacked *traX* and *traY*.

### 3.5. pOXA-48 Variant 3

pOXA-48 variant 3 shares a similar structure to the first variant. Variant 3 however encodes only 4 AMR genes, *bla*_OXA-48_, *bla*_CTX-M-14b_, *aph(3’’)-Ib* and *aph(6)-Id*. Both *aph(3’’)-Ib* and *aph(6)-Id*, encoding aminoglycoside 3’-phosphotransferase and aminoglycoside O-phosphotransferase, in addition to the resolvase bracketed between IS26 and the ISEc63 insertion transposases, forming a 6009 bp fragment. Similar to pOXA-48 variant 1, variant 3 has a structure similar to modules belonging to the transposon, Tn5393, with a 100% nucleotide match for the genes *aph(3”)-Ib* and, *aph(6)-Id*, and a 99% identity for the resolvase. Together, this structure may represent a potential composite transposon. The IS26-ISEc63 AMR fragment is inserted between a hypothetical gene downstream of *ymoA* and intergenic region upstream of the *klcA* gene. The ISEcp1-*bla*_CTX-M-14b_ fragment is inserted downstream of the z1226 gene in the intergenic region. The 2 insertions relative to the reference pOXA-48 plasmid is shown in Figure 7 (top). The positioning of the IS26-ISEc63 fragment with the AMR genes in linkage, and the ISEcp1-*bla*_CTX-M-14b_ is shown for representative plasmid pSA-KpST14-OXA48-2 in Figure 7.

**Figure 7.**
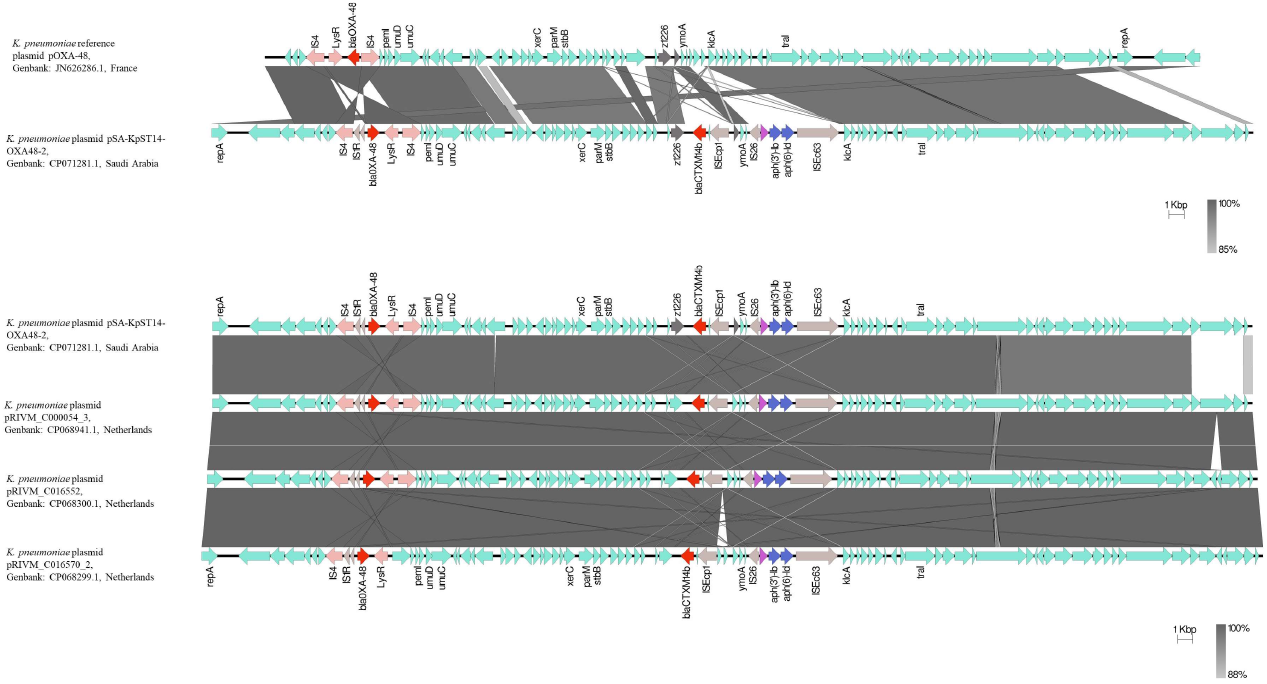
AMR insertions in the pOXA-48 variant 3 plasmid. Top. On representative plasmid pSA-KpST14-OXA48-2, the fragment is inserted between 73 nucleotides downstream of a hypothetical gene downstream of *ymoA*, and the stop codon of a hypothetical gene in the intergenic region upstream of the *klcA* gene. The ISEcp1-_*bla*CTX-M-14b_ fragment is inserted 37 nucleotides downstream of the *z1226* gene in the intergenic region. Bottom. Both the ISEcp1-_*bla*CTX-M-14b_ and the IS26-ISEc63 AMR insertions positioned on the pOXA-48 plasmid. Antibiotic resistance genes identified using ResFinder v.4.0 (Zankari *et al*., 2012).

The BLASTn program from NCBI revealed the presence of the 6009 bp IS26-ISEc63 AMR fragment in a total of 6 plasmid sample; 5 plasmid samples from the pOXA-48 samples (Figure 2b; Table 3; Supplementary Table S1), and an additional plasmid sample, *E.coli* plasmid, pRCS60_p submitted from the Genoscope sequencing center in France. While the IS26-ISEc63 fragment was identified in 6 plasmid samples altogether, only 5 pOXA-48-like plasmids harbored the complete IS26-ISEc63 fragment. Plasmid sample pRIVM_C018614_3 had truncations in the genes encoding *aph(3”)-Ib* and, *aph(6)-Id* and the ISEc63 transposase. The plasmid sample from France, pRCS60_p lacked the *bla*_OXA-48_ gene. All 6 samples did encode the *bla*_CTX-M-14b_ gene. Among the plasmids from the Netherlands, the 3 plasmids were between 1-46 SNPs distant from plasmid pRIVM_C000054_3, indicating high similarity between the 4 plasmids. In fact, the plasmids from the Netherlands had ≥ 99.87% identity and 100% coverage against pRIVM_C000054_3 (Table 3). The two plasmid samples from the Saudi Arabia, and France, pSA-KpST14 and pRCS60_p, were however, 645 and 678 SNPs distant from plasmid pRIVM_C000054_3 from the Netherlands, displaying high identity, ≥99.44%, but lower coverage at 94%. Interestingly, both the both the plasmids from Saudi Arabi and France are only 37 SNPs distant from one another.

**Table 3.**
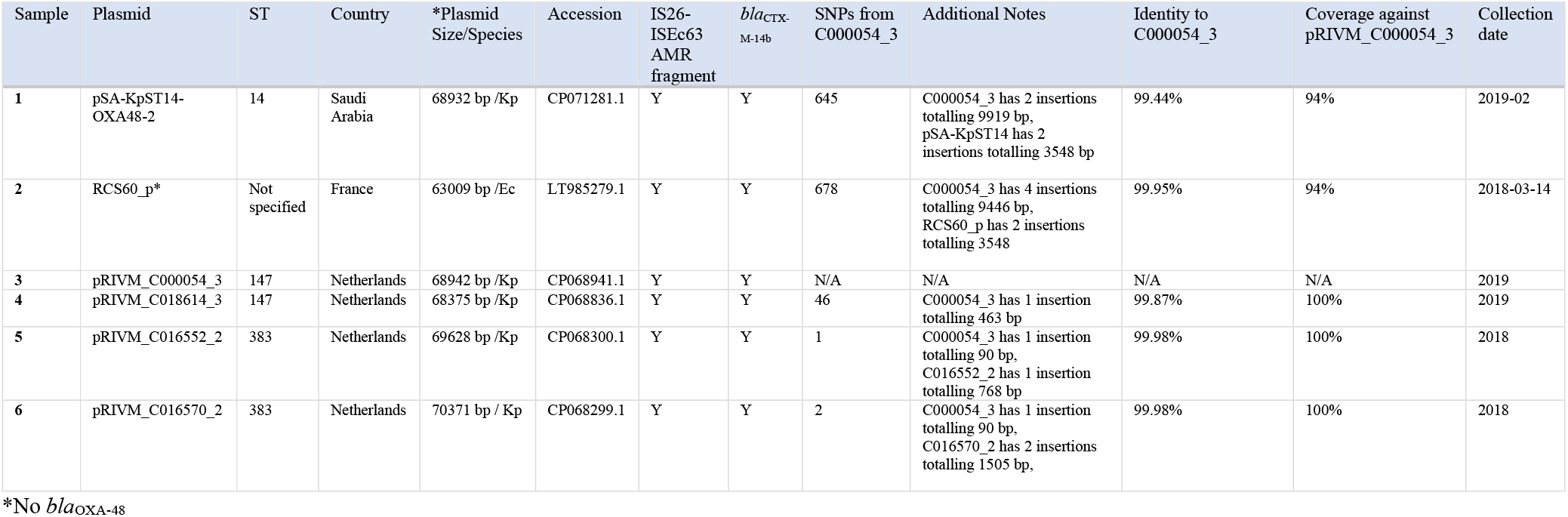
pOXA-48 variant 3 relatedness.

| Sample | Plasmid | ST | Country | *Plasmid Size/Species | Accession | IS26-ISEc63 AMR fragment | *bla*<sub>CTX-M-14b</sub> | SNPs from C000054_3 | Additional Notes | Identity to C000054_3 | Coverage against pRIVM_C000054_3 | Collection date |
| --- | --- | --- | --- | --- | --- | --- | --- | --- | --- | --- | --- | --- |
| 1 | pSA-KpST14-OXA48-2 | 14 | Saudi Arabia | 68932 bp /Kp | CP071281.1 | Y | Y | 645 | C000054_3 has 2 insertions totalling 9919 bp, pSA-KpST14 has 2 insertions totalling 3548 bp | 99.44% | 94% | 2019-02 |
| 2 | RCS60_p* | Not specified | France | 63009 bp /Ec | LT985279.1 | Y | Y | 678 | C000054_3 has 4 insertions totalling 9446 bp, RCS60 p has 2 insertions totalling 3548 | 99.95% | 94% | 2018-03-14 |
| 3 | pRIVM_C000054_3 | 147 | Netherlands | 68942 bp /Kp | CP068941.1 | Y | Y | N/A | N/A | N/A | N/A | 2019 |
| 4 | pRIVM_C018614_3 | 147 | Netherlands | 68375 bp /Kp | CP068836.1 | Y | Y | 46 | C000054_3 has 1 insertion totalling 463 bp | 99.87% | 100% | 2019 |
| 5 | pRIVM_C016552_2 | 383 | Netherlands | 69628 bp /Kp | CP068300.1 | Y | Y | 1 | C000054_3 has 1 insertion totalling 90 bp, C016552_2 has 1 insertion totalling 768 bp | 99.98% | 100% | 2018 |
| 6 | pRIVM_C016570_2 | 383 | Netherlands | 70371 bp / Kp | CP068299.1 | Y | Y | 2 | C000054_3 has 1 insertion totalling 90 bp, C016570_2 has 2 insertions totalling 1505 bp, | 99.98% | 100% | 2018 |

\*No *bla*<sub>OXA-48</sub>

#### 3.5.1 *Tra* Transfer Operon for pOXA-48 variant 3

Among the plasmid samples from the Netherlands, the 4 plasmid samples had a 100% coverage and 99.85% percentage identity against the *tra* operon, HIJK*pri*LMNOPQRUWXY. The similar coverage and identity reflect the high similarity between the plasmids. The two plasmid samples from Saudi Arabi and France, both had 85% coverage and 97.6% nucleotide identity against the *tra* operon, whereby both plasmid samples did not encode either the *traX* or *traY* genes, respectively. Collectively, these results infer all pOXA-48 variant 3 plasmids are likely conjugative.

## 4. Discussion

The *bla*_OXA-48_ allele reveal the gene was largely limited to IncL/M pOXA-48 like plasmids harboring only *bla*_OXA-48_. Concerningly, 4 plasmids encoding the *bla*_OXA-48_ gene were sourced from human food consumption samples. The presence of these carbapenemase genes among food samples provides a route for pOXA-48 plasmid acquisition in community settings. Furthermore, the BLASTn hits revealed the presence of 34 pOXA-48 like plasmids hits carrying additional AMR genes beyond *bla*_OXA-48_. The presence of AMR genes on pOXA-48 plasmids was associated with samples sourced from clinical/surveillance settings. This may indicate antibiotic pressure in hospitals promotes the generation of MDR strains.

*In-silico* antibiotic resistance gene analyses identified three common variants of the pOXA-48 plasmid encoding aminoglycoside, *bla*_CTX-M-14b_ and *bla*_OXA-48_ resistance genes. Notably, pOXA-48 variant 1 encoding *aph(3”)-Ib* (2), *aph(6)-Id, aph(3’)-Vib*, *bla*_CTX-M-14b_, and *bla*_OXA-48_ AMR genes were identified in 7 unique countries including, including Germany (*n*=2), China (*n*=6), UK (*n*=1), USA (*n*=1), Palestine (*n*=1), Australia (*n*=1), and the Netherlands (*n*=2). 13 plasmids carrying the 6 AMR genes common to pOXA-48 variant 1 were closely related to pOXA48-IR1251 (accession: CP068574.1), from China. 13 plasmids had a nucleotide identity ≥ 99.29% and a coverage of ≥92%, indicating their potential international dissemination. Further, genomic surveillance of pOXA-48 like plasmids will be required to establish their persistence more fully. pOXA-variant 3 appears more confined internationally, although a similar plasmid was detected in multiple samples from the Netherlands. pOXA variant 2 included highly similar plasmids present in three independent countries, suggesting either clonal expansion of isolates with this plasmid or HGT promotes dissemination.

Worryingly, 4 patients infected with a *K. pneumoniae* isolate from an ICU unit in Australia harboring the pJEG011 plasmid, carrying aminoglycoside/*bla*_CTX-M-14_, and *bla*_OXA-48_ AMR genes died (Espedido *et al*., 2013), while another OXA-48 producing *K. pneumoniae* co-producing a similar ESBL, CTX-M-15, caused a nosocomial outbreak resulting in patients’ deaths (Pantel *et al*., 2016). Conjugations experiments revealed pJEG011 can be readily transferred from the original *K. pneumoniae* isolate to recipient *E.coli* cells. While transconjugants *E.coli* cells displayed an increased minimal inhibitory concentration (MIC) towards β-lactams, the transconjugant remained susceptible to meropenem (Espedido *et al*., 2013). Notably, the donor *K. pneumoniae* isolate had mutational insertions in outer membrane porins, OmpK35 and OmpK36, proteins necessary for carbapenem entry into the cell. Carbapenem resistance in *bla*_OXA-48_ containing isolates is usually co-dependent on additional resistance mechanisms, such as mutations in outer membrane porins, OmpK35 and OmpK36. All pOXA-48 AMR plasmid variants may have the capacity to go undetected in clinical settings based on antibiotic susceptibility profiles promoting dissemination of ESBL and carbapenem resistance genes. pOXA-variants 1 and 3 have a similar genomic architecture. Notably, the aminoglycoside resistance genes were bracketed between 2 insertion sequence elements, IS26 and ISEc63, encoding transposases. In between these IS elements, a Tn3 resolvase was also found. Combined, this structure may form a composite transposon. The potential transposon structure had resemblance to the Tn5393 transposon (accession: M96392), including both *aph(3’’)-Ib, aph(6)-Id* genes, and a Tn3 resolvase. Across both pOXA-48 variants 1 and 3, this complete transposon structure was identified in 9 separate countries on plasmids both closely related and more distantly related suggesting both HGT and independent import into pOXA-48 may promote acquisition of the potential transposon in recipient strains. Crucially, this may mean, multiple AMR genes can shuttle into pOXA-48 IncL plasmids in a single step, expanding the resistome of recipient strains. Future functional studies exploring using both pOXA-48 variants are necessary to prove this mechanism. For all pOXA-48 variant 1 and 3 samples, the IS element, ISEcp1 was located upstream of the ESBL gene *bla*_CTX-M-14b_. ISEcp1 upstream of *CTX-M* genes is a common genomic configuration reported (Khalaf *et al*., 2009; Tian *et al*., 2011; Shawa *et al*., 2021), leading to the speculation that ISEcp1 may be responsible for *bla*_CTX-M_ transposition, and the development of multi-drug resistant strains.

All plasmid AMR variants encoded one of two variants of the *tra* operon, HIJKpriLMNOPQRUWXY encoding conjugative transfer machinery. Mutation and conjugation assays confirm the *tra* operon with and without *traXY* maintain DNA conjugative transfer ability (Maneewannakul *et al*., 1995; Lang *et al*., 2012; Chen *et al*., 2015) indicating all plasmids may maintain the ability to transfer AMR genes co-residing on the same plasmid.

pOXA-48 dissemination between patients is mediated by the spread of high-risk *K. pneumoniae* clones, such as ST11 (León-Sampedro *et al*., 2021). Multiple high-risk *K. pneumoniae* STs carrying pOXA-48 AMR plasmid variants were detected, including ST11, 101, 147, and 383. Among colonized patients, pOXA-48 readily spreads via conjugation to other resident members of the gut microbiota. Conjugation can lead to the long-term establishment of pOXA-48 mediated by HGT of pOXA-48. The presence of pOXA-48 variants harboring additional AMR genes beyond *bla*_OXA-48_ in high-risk clones may lead to similar between-and-within patient transfer dynamics. Lengthier colonization periods in clinical settings may provide more opportunities for between patient transfer.

The presence of the AMR genes common to pOXA-48 variants 1/2/3 may severely limit therapeutic treatment options. An *E.coli* transconjugant carrying *bla*_OXA-48_ on a pOXA-48 plasmid degraded the broad-spectrum cephalosporin cefotaxime into two inactive compounds, indicating cefotaxime is not a valid therapeutic for treating infections caused by *bla*_OXA-48_-producing bacteria, rather, ceftazidime and ceftriaxone remain valid choices (Oviaño *et al*., 2019). The presence of the *bla*_CTX-M-14_/*bla*_CTX-M-14b_ on plasmids in *E.coli*, however, has been associated with MICs >= 16 mg/L for the third-generation cephalosporins, cefotaxime, and ceftazidime (Brinas *et al*., 2005; Liao *et al*., 2015). Furthermore, conjugation of the plasmid pML2508 encoding *bla*_CTX-M-14-b_ caused an increase in the MIC of cefotaxime, ceftazidime, ceftriaxone, and cefpirome from 0.13 μg/ml to 64 μg/ml for ceftazidime and >512 μg/ml for the other cephalosporins (Achour *et al*., 2012). Taken together, the co-occurrence of *bla*_OXA-48_ and *bla*_CTX-M-14-b_ severely limits treatment options with both carbapenems and cephalosporins.

Aminoglycoside antibiotics including amikacin, gentamicin, kanamycin, streptomycin, and tobramycin alongside carbapenems such as meropenem/imipenem represent valid combination treatment therapies yielding synergistic effects for infections caused by carbapenem resistant Enterobacteriaceae (CRE) (Hajjej *et al*., 2016; Terbtothakun *et al*., 2021). The presence of multiple aminoglycoside genes encoding aminoglycoside-modifying enzymes (AMEs) among all three pOXA-48 AMR variants may prevent synergistic therapies involving both carbapenems and aminoglycosides.

Analysis was limited to sequenced plasmids sourced from NCBI. The *bla*_OXA-48_ gene is present in species not recorded in our dataset, including *Pseudomonas aeruginosa* whereby PCR assays confirm the presence of the carbapenemase gene (Ali and Nagla, 2020). Sourcing plasmids from NCBI may, if anything underrepresent the complete epidemiology of pOXA-48 like plasmids and hence the scale of the problem. The discovery of aminoglycoside, ESBL, and carbapenemase resistance genes in a single broad host conjugative plasmid in clinical settings is particularly worrying.

Another limitation of the study was AMR genes were only detected using sequenced non-duplicate plasmids. Antimicrobial susceptibility testing (AST) data was not available to confirm the predicted AMR phenotype. To help mitigate against this effect, the latest version of ResFinder v.4.0 was used, which has shown high genotype–phenotype concordance for the Gram-negative bacterium *E.coli* for the prediction of resistance towards meropenem (100%), cefepime (100%), ciprofloxacin (99.2%), and gentamicin (97.6%) using human and animal clinical samples and surveillance samples (Bortolaia *et al*., 2020). Importantly, the AMR genes detected in the pOXA-48 samples confer resistance towards these clinically relevant antibiotics.

Sequenced plasmid contigs reveal the presence of pOXA-48 variants carrying multiple AMR genes beyond *bla*_OXA-48_. Acquisition and carriage of pOXA-48 resistant plasmids can change the resistome of susceptible isolates in a single-step process, rendering previously susceptible strains refractory to almost all available treatment options.

## Appendix

**Table A1.**
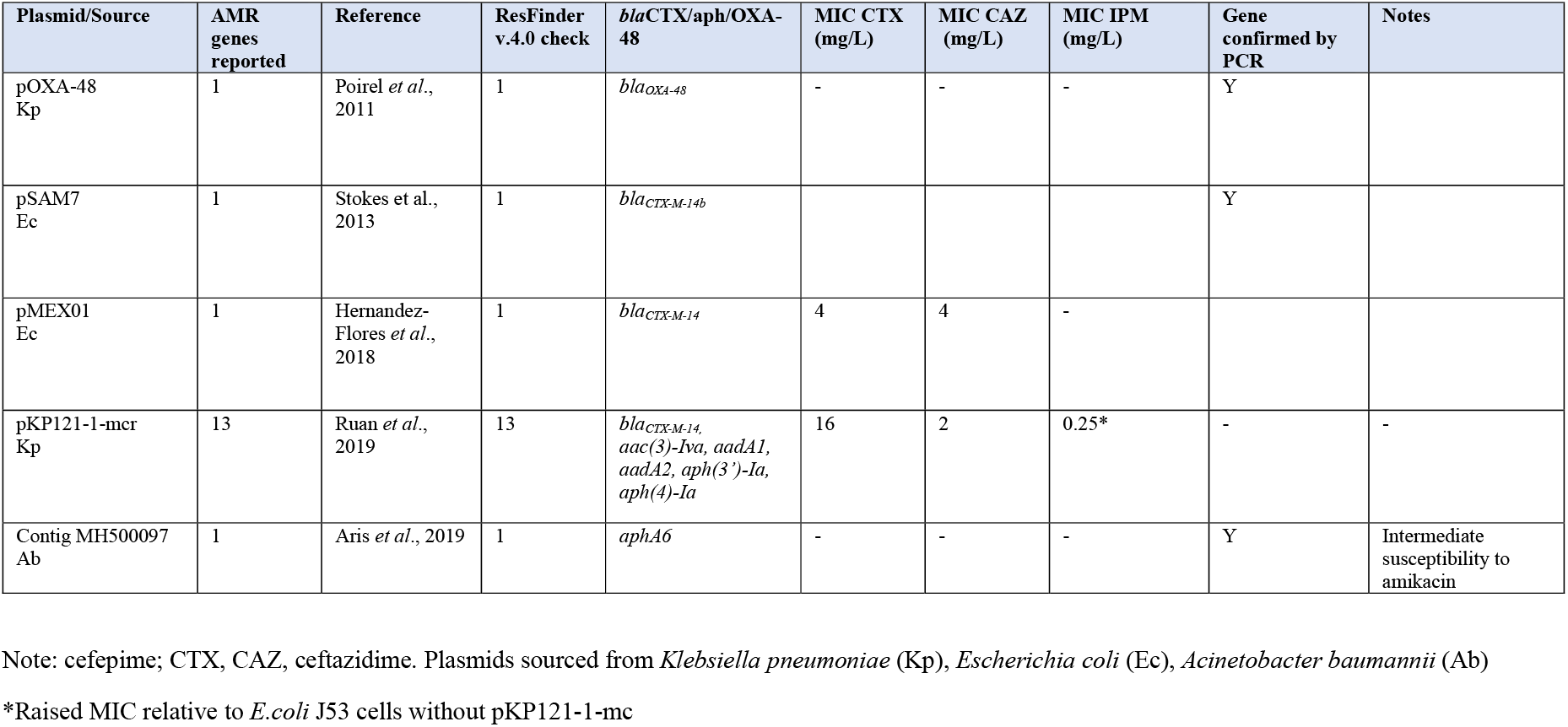
Comparison of confirmed AMR genes using PCR/MIC values and predictions from ResFinder v.4.0.

## References

1. Ali, D. O., & Nagla, M. M. (2020). Molecular detection of BlaOXA-48 gene encoding Carbapenem resistance pseudomonas aeruginosa clinical isolates from Khartoum state hospitals, Sudan. Gezira Journal of Health Sciences, 17, https://doi.org/10.1101/2020.06.22.20137034

2. Aris, P., Boroumand, M. A., & Douraghi, M. (2019). Amikacin resistance due to the aphA6 gene in multi-antibiotic resistant *Acinetobacter baumannii* isolates belonging to global clone 1 from Iran. BMC Microbiology, 19(1). https://doi.org/10.1186/s12866-019-1592-6

3. Balkan, I. I., Aygün, G., Aydın, S., Mutcalı, S. I., Kara, Z., Kuşkucu, M., Midilli, K., Şemen, V., Aras, Ş., Yemişen, M., Mete, B., Özaras, R., Saltoğlu, N., Tabak, F., & Öztürk, R. (2014). Blood stream infections due to OXA-48-like carbapenemase-producing *Enterobacteriaceae*: Treatment and survival. International Journal of Infectious Diseases, 26, 51–56. https://doi.org/10.1016/j.ijid.2014.05.012

4. Ben Achour, N., Power, P., Mercuri, P. S., Ben Moussa, M., Moreno, G., & Belhadj, O. (2012). First detection of a transferable *bla*_CTX-M-14b_ gene in a *Klebsiella pneumoniae* clinical isolate from Tunisia and analysis of its genetic context. Annals of Microbiology, 62(4), 1737–1742. https://doi.org/10.1007/s13213-012-0430-y

5. Bortolaia, V., Kass, R. S., Ruppe, E., Roberts, M. C., Schwarz, S., Cattoir, V., Philippon, A., Allesoe, R. L., Rebelo, A. R., Florensa, A. F., Fagelhauer, L., Chakraborty, T., Neumann, B., Werner, G., Bender, J. K., Stingl, K., Nguyen, M., Coppens, J., Xavier, B. B., Malhotra-Kumar, S., Westh, H., Pinholt, M., Anjum, M. F., Duggett, N. A., Kempf, I., Nykasenoja, S., Olkkola, S., Wieczorek, K., Amaro, A., Clemente, L., Mossong, J., Losch, S., Ragimbeau, C., Lund, O., Aarestrup, M. A. (2020). ResFinder 4.0 for predictions of phenotypes from genotypes. Journal of Antimicrobial Chemotherapy, 75, 3491–3500. https://doi.org/10.1093/jac/dkaa345

6. Briñas, L., Lantero, M., De Diego, I., Alvarez, M., Zarazaga, M., & Torres, C. (2005). Mechanisms of resistance to expanded-spectrum cephalosporins in *Escherichia coli* isolates recovered in a Spanish hospital. Journal of Antimicrobial Chemotherapy, 56(6), 1107–1110. https://doi.org/10.1093/jac/dki370

7. Carattoli, A., & Hasman, H. (2019). PlasmidFinder and in Silico pMLST: Identification and typing of plasmid replicons in whole-genome sequencing (WGS). Horizontal Gene Transfer, 285–294. https://doi.org/10.1007/978-1-4939-9877-7_20

8. Carver, T. J., Rutherford, K. M., Berriman, M., Rajandream, M., Barrell, B. G., & Parkhill, J. (2005). ACT: The Artemis comparison tool. Bioinformatics, 21(16), 3422–3423. https://doi.org/10.1093/bioinformatics/bti553

9. Chen, L., Al Laham, N., Chavda, K. D., Mediavilla, J. R., Jacobs, M. R., Bonomo, R. A., & Kreiswirth, B. N. (2015). First report of an OXA-48-Producing multidrug-resistant *Proteus mirabilis* strain from Gaza, Palestine. Antimicrobial Agents and Chemotherapy, 59(7), 4305–4307. https://doi.org/10.1128/aac.00565-15

10. Chen, L., Al Laham, N., Chavda, K. D., Mediavilla, J. R., Jacobs, M. R., Bonomo, R. A., & Kreiswirth, B. N. (2015). First report of an OXA-48-Producing multidrug-resistant *Proteus mirabilis* strain from Gaza, Palestine. Antimicrobial Agents and Chemotherapy, 59(7), 4305–4307. https://doi.org/10.1128/aac.00565-15

11. Dimou, V., Dhanji, H., Pike, R., Livermore, D. M., & Woodford, N. (2012). Characterization of Enterobacteriaceae producing OXA-48-like carbapenemases in the UK. Journal of Antimicrobial Chemotherapy, 67(7), 1660–1665. https://doi.org/10.1093/jac/dks124

12. Espedido, B. A., Steen, J. A., Ziochos, H., Grimmond, S. M., Cooper, M. A., Gosbell, I. B., Van Hal, S. J., & Jensen, S. O. (2013). Whole genome sequence analysis of the first Australian OXA-48-Producing outbreak-associated *Klebsiella pneumoniae* isolates: The Resistome and in vivo evolution. PLoS ONE, 8(3), e59920. https://doi.org/10.1371/journal.pone.0059920

13. Fei Tian, S., Zhuo Chu, Y., Yi Chen, B., Nian, H., & Shang, H. (2011). ISEcp1 element in association with *bla*_CTX-M_ genes of *E. coli* that produce extended-spectrum β-lactamase among the elderly in community settings. Enfermedades Infecciosas y Microbiología Clínica, 29(10), 731–734. https://doi.org/10.1016/j.eimc.2011.07.011

14. Findlay, J., Hopkins, K. L., Loy, R., Doumith, M., Meunier, D., Hill, R., Pike, R., Mustafa, N., Livermore, D. M., & Woodford, N. (2017). OXA-48-like carbapenemases in the UK: An analysis of isolates and cases from 2007 to 2014. Journal of Antimicrobial Chemotherapy, 72(5), 1340–1349. https://doi.org/10.1093/jac/dkx012

15. Hajjej, Z., Gharsallah, H., Naija, H., Boutiba, I., Labbene, I., & Ferjani, M. (2016). Successful treatment of a carbapenem-resistant *Klebsiella pneumoniae* carrying *bla*_OXA-48_, *bla*_VIM-2_, *bla*_CMY-2_ and *bla*_SHV_-with high dose combination of imipenem and amikacin. IDCases, 4, 10–12. https://doi.org/10.1016/j.idcr.2016.01.003

16. Hamprecht, A., Sommer, J., Willmann, M., Brender, C., Stelzer, Y., Krause, F. F., Tsvetkov, T., Wild, F., Riedel-Christ, S., Kutschenreuter, J., Imirzalioglu, C., Gonzaga, A., Nübel, U., & Göttig, S. (2019). Pathogenicity of clinical OXA-48 isolates and impact of the OXA-48 IncL plasmid on virulence and bacterial fitness. Frontiers in Microbiology, 10. https://doi.org/10.3389/fmicb.2019.02509

17. Hendrickx, A. P., Landman, F., De Haan, A., Borst, D., Witteveen, S., Van Santen-Verheuvel, M., & Schouls, L. M. (2020). *bla*OXA-48-like genome architecture among carbapenemase-producing *Escherichia coli* and *Klebsiella pneumoniae* in The Netherlands. Microbial Genomics, 7(5). https://doi.org/10.1099/mgen.0.000512

18. Hernandez-Flores, J. L., Pérez, J. C., Gutiérrez, C. S., Hernández, A. C., Alonso, G. S., Hernández, S. P., Gómez, S. R., Fernández, F., Loske, A. M., & Guillén, J. C. (2018). PMEX01, a 70 kb plasmid isolated from *Escherichia coli* that confers resistance to multiple β-lactam antibiotics. Brazilian Journal of Microbiology, 49(3), 569–574. https://doi.org/10.1016/j.bjm.2017.11.002

19. Hernández-García, M., Pérez-Viso, B., Navarro-San Francisco, C., Baquero, F., Morosini, M. I., Ruiz-Garbajosa, P., & Cantón, R. (2019). Intestinal Co-Colonization with different carbapenemase-producing Enterobacterales isolates is not a rare event in an OXA-48 endemic area. EClinicalMedicine, 15, 72–79. https://doi.org/10.1016/j.eclinm.2019.09.005

20. Khalaf, N. G., Eletreby, M. M., & Hanson, N. D. (2009). Characterization of CTX-M ESBLs in *Enterobacter cloacae*, *Escherichia coli* and *Klebsiella pneumoniae* clinical isolates from Cairo, Egypt. BMC Infectious Diseases, 9(1). https://doi.org/10.1186/1471-2334-9-84

21. Lang, S., & Zechner, E. L. (2012). General requirements for protein secretion by the F-like conjugation system R1. Plasmid, 67(2), 128–138. https://doi.org/10.1016/j.plasmid.2011.12.014

22. Lang, S., & Zechner, E. L. (2012). General requirements for protein secretion by the F-like conjugation system R1. Plasmid, 67(2), 128–138. https://doi.org/10.1016/j.plasmid.2011.12.014

23. León-Sampedro, R., DelaFuente, J., Díaz-Agero, C., Crellen, T., Musicha, P., Rodríguez-Beltrán, J., De la Vega, C., Hernández-García, M., López-Fresneña, N., Ruiz-Garbajosa, P., Cantón, R., Cooper, B. S., & San Millán, Á. (2021). Pervasive transmission of a carbapenem resistance plasmid in the gut microbiota of hospitalized patients. Nature Microbiology, 6(5), 606–616. https://doi.org/10.1038/s41564-021-00879-y

24. Liao, X., Xia, J., Yang, L., Li, L., Sun, J., Liu, Y., & Jiang, H. (2015). Characterization of CTX-M-14-producing *Escherichia coli* from food-producing animals. Frontiers in Microbiology, 6. https://doi.org/10.3389/fmicb.2015.01136

25. Lim, F., Modha, D., Collins, E., Westmoreland, D., Ashton, C., & Jenkins, D. (2020). An outbreak of two strains of OXA-48 producing *Klebsiella pneumoniae* in a teaching hospital. Infection Prevention in Practice, 2(3), 100033. https://doi.org/10.1016/j.infpip.2019.100033

26. Madueño, A., González-García, J., Alonso Socas, M. D., Miguel Gómez, M. A., & Lecuona, M. (2018). Clinical features and outcomes of bacteraemia due to OXA-48-like carbapenemase-producing *Klebsiella pneumoniae* in a tertiary hospital. Enfermedades infecciosas y microbiologia clinica (English ed.), 36(8), 498–501.

27. Maneewannakul, K., Maneewannakul, S., & Ippen-Ihler, K. (1995). Characterization of traX, the F plasmid locus required for acetylation of F-pilin subunits. Journal of Bacteriology, 177(11), 2957–2964. https://doi.org/10.1128/jb.177.11.2957-2964.1995

28. Navarro-San Francisco, C., Mora-Rillo, M., Romero-Gómez, M., Moreno-Ramos, F., Rico-Nieto, A., Ruiz-Carrascoso, G., Gómez-Gil, R., Arribas-López, J., Mingorance, J., & Paño-Pardo, J. (2013). Bacteraemia due to OXA-48-carbapenemase-producing Enterobacteriaceae: A major clinical challenge. Clinical Microbiology and Infection, 19(2), E72–E79. https://doi.org/10.1111/1469-0691.12091

29. Oviaño, M., Rodicio, M. R., Heinisch, J. J., Rodicio, R., Bou, G., & Fernández, J. (2019). Analysis of the degradation of broad-spectrum cephalosporins by OXA-48-Producing Enterobacteriaceae using MALDI-TOF MS. Microorganisms, 7(12), 614. https://doi.org/10.3390/microorganisms7120614

30. Page, A. J., Cummins, C. A., Hunt, M., Wong, V. K., Reuter, S., Holden, M. T., Fookes, M., Falush, D., Keane, J. A., & Parkhill, J. (2015). Roary: Rapid large-scale prokaryote pan genome analysis. Bioinformatics, 31(22), 3691–3693. https://doi.org/10.1093/bioinformatics/btv421

31. Poirel, L., Bonnin, R. A., & Nordmann, P. (2011). Genetic features of the widespread plasmid coding for the Carbapenemase OXA-48. Antimicrobial Agents and Chemotherapy, 56(1), 559–562. https://doi.org/10.1128/aac.05289-11

32. Poirel, L., Héritier, C., Tolün, V., & Nordmann, P. (2004). Emergence of oxacillinase-mediated resistance to Imipenem in klebsiella pneumoniae. Antimicrobial Agents and Chemotherapy, 48(1), 15–22. https://doi.org/10.1128/aac.48.1.15-22.2004

33. Potron, A., Nordmann, P., Rondinaud, E., Jaureguy, F., & Poirel, L. (2012). A mosaic transposon encoding OXA-48 and CTX-M-15: Towards pan-resistance. Journal of Antimicrobial Chemotherapy, 68(2), 476–477. https://doi.org/10.1093/jac/dks397

34. Public Health England. Detection of bacteria with carbapenem hydrolysing β lactamases (carbapenemases). (2016, September 16). GOV.UK. https://www.gov.uk/government/publications/smi-b-60-detection-of-bacteria-with-carbapenem-hydrolysing-lactamases-carbapenemases

35. Rodríguez, O. L., Sousa, A., Pérez-Rodríguez, M. T., Martínez-Lamas, L., Suárez, R. L., Martínez, C. T., Pino, C. P., Vidal, F. V., Pérez-Landeiro, A., & Casal, M. C. (2021). Mortality-related factors in patients with OXA-48 carbapenemase-producing *Klebsiella pneumoniae* bacteremia. Medicine, 100(14), e24880. https://doi.org/10.1097/md.0000000000024880

36. Ruan, Z., Sun, Q., Jia, H., Huang, C., Zhou, W., Xie, X., & Zhang, J. (2019). Emergence of a ST2570 *Klebsiella pneumoniae* isolate carrying mcr-1 and *bla*_CTX-M-14_ recovered from a bloodstream infection in China. Clinical Microbiology and Infection, 25(7), 916–918. https://doi.org/10.1016/j.cmi.2019.02.005

37. Shaidullina, E., Shelenkov, A., Yanushevich, Y., Mikhaylova, Y., Shagin, D., Alexandrova, I., Ershova, O., Akimkin, V., Kozlov, R., & Edelstein, M. (2020). Antimicrobial resistance and Genomic characterization of OXA-48-and CTX-M-15-Co-Producing Hypervirulent *Klebsiella pneumoniae* ST23 recovered from nosocomial outbreak. Antibiotics, 9(12), 862. https://doi.org/10.3390/antibiotics9120862

38. Shawa, M., Furuta, Y., Mulenga, G., Mubanga, M., Mulenga, E., Zorigt, T., Kaile, C., Simbotwe, M., Paudel, A., Hang’ombe, B., & Higashi, H. (2021). Novel chromosomal insertions of ISEcp1-*bla*_CTX-M-15_ and diverse antimicrobial resistance genes in Zambian clinical isolates of *Enterobacter cloacae* and *Escherichia coli*. Antimicrobial Resistance & Infection Control, 10(1). https://doi.org/10.1186/s13756-021-00941-8

39. Stokes, M. O., AbuOun, M., Umur, S., Wu, G., Partridge, S. R., Mevius, D. J., Coldham, N. G., & Fielder, M. D. (2013). Complete sequence of pSAM7, an IncX4 plasmid carrying a Novel *bla*CTX-M-14bTransposition unit isolated from *Escherichia coli* and *Enterobacter cloacae* from cattle. Antimicrobial Agents and Chemotherapy, 57(9), 4590–4594. https://doi.org/10.1128/aac.01157-13

40. Sullivan, M. J., Petty, N. K., & Beatson, S. A. (2011). Easyfig: A genome comparison visualizer. Bioinformatics, 27(7), 1009–1010. https://doi.org/10.1093/bioinformatics/btr039

41. Taoufik, L., Amrani Hanchi, A., Fatiha, B., Nissrine, S., Mrabih Rabou, M. F., & Nabila, S. (2019). Emergence of OXA-48 Carbapenemase producing *Klebsiella pneumoniae* in a neonatal intensive care unit in Marrakech, Morocco. Clinical Medicine Insights: Pediatrics, 13, 117955651983452. https://doi.org/10.1177/1179556519834524

42. Terbtothakun, P., Nwabor, O. F., Siriyong, T., Voravuthikunchai, S. P., & Chusri, S. (2021). Synergistic antibacterial effects of Meropenem in combination with Aminoglycosides against carbapenem-resistant *Escherichia coli* harboring *bla*_NDM-1_ and *bla*_NDM-5_. Antibiotics, 10(8), 1023. https://doi.org/10.3390/antibiotics10081023

43. Valle, A. A., León-Sampedro, R., Rodríguez-Beltrán, J., DelaFuente, J., Hernández-García, M., Ruiz-Garbajosa, P., Cantón, R., Peña-Miller, R., & Millán, Á. S. (2020). The distribution of plasmid fitness effects explains plasmid persistence in bacterial communities. Nature Communications 12(2653) https://doi.org/10.1101/2020.08.01.230672

44. Wang, L., Guo, L., Ye, K., & Yang, J. (2021). Genetic characteristics of OXA-48-producing Enterobacterales from China. Journal of Global Antimicrobial Resistance, 26, 285–291. https://doi.org/10.1016/j.jgar.2021.07.006

45. Woodford, N., Eastaway, A. T., Ford, M., Leanord, A., Keane, C., Quayle, R. M., Steer, J. A., Zhang, J., & Livermore, D. M. (2010). Comparison of BD Phoenix, Vitek 2, and MicroScan automated systems for detection and inference of mechanisms responsible for Carbapenem resistance in Enterobacteriaceae. Journal of Clinical Microbiology, 48(8), 2999–3002. https://doi.org/10.1128/jcm.00341-10

46. Zankari, E., Hasman, H., Cosentino, S., Vestergaard, M., Rasmussen, S., Lund, O., Aarestrup, F. M., & Larsen, M. V. (2012). Identification of acquired antimicrobial resistance genes. Journal of Antimicrobial Chemotherapy, 67(11), 2640–2644. https://doi.org/10.1093/jac/dks261

